# Eco-evolutionary community turnover following environmental change

**DOI:** 10.1101/285288

**Authors:** Jesse R. Lasky

## Abstract

Co-occurring species often differ in intraspecific genetic diversity, which in turn can affect adaptation in response to environmental change. Specifically, the simultaneous evolutionary responses of co-occurring species to temporal environmental change may influence community dynamics. Local adaptation along environmental gradients combined with gene flow can enhance genetic diversity of traits within populations. Quantitative genetic theory shows that having greater gene flow results in (1) lower equilibrium population size due to maladaptive immigrant genotypes (migration load), but (2) faster adaptation to changing environments. Here I build off this theory to study community dynamics of locally adapted species in response to temporal environmental changes akin to warming temperatures. Although an abrupt environmental change leaves all species initially maladapted, high gene flow species subsequently adapt faster due to greater genetic diversity. As a result, species can transiently reverse their relative abundances, but sometimes only after long lag periods. If constant temporal environmental change is applied, the community exhibits a shift toward stable dominance by species with intermediate gene flow. Notably, fast-adapting high gene flow species can increase in absolute abundance under environmental change (although often only for a transient period) because the change suppresses superior competitors with lower gene flow. This eco-evolutionary competitive release stabilizes ecosystem function. The eco-evolutionary community turnover studied here parallels the purely ecological successional dynamics following disturbances. My results demonstrate how interspecific variation in life history can have far-reaching impacts on eco-evolutionary community response to environmental change.

## Introduction

Genetic diversity in quantitative traits serves as the raw material for selection (Lush 1937). Understanding how rapid changes in selection impact populations is a question with tremendous importance in biodiversity conservation, agriculture, and medicine (Gomulkiewicz and Holt 1995, Bell and Gonzalez 2009, Read et al. 2011, Alexander et al. 2014, Lasky et al. 2015, Bay et al. 2017). A substantial portion of genetic diversity in phenotypes within species is maintained due to population adaptation to local environments (Turesson 1922, Clausen et al. 1940, Leimu and Fischer 2008, Hereford 2009). Local adaptation is defined as a genotype-by-environment interaction favoring home genotypes (Kawecki and Ebert 2004). When populations are locally adapted, greater gene flow can increase within-population diversity due to immigration from populations adapted to other environments (Barton 2001, Lenormand 2002, Garant et al. 2007). Given that local adaptation is common (Leimu and Fischer 2008, Hereford 2009, Sanford and Kelly 2010) and multiple co-occurring species can be simultaneously adapted to local environments, these processes could impact genetic diversity of co-occurring species and community responses to environmental change. Here I build on previous theory to study the complex role gene flow plays in communities due to its effect on genetic diversity, which induces migration load on populations but also speeds up adaptation (Pease et al. 1989, Polechová et al. 2009, Kremer et al. 2012).

A major body of theory explores the conditions under which selective gradients lead to stable polymorphism and local adaptation (Haldane 1930, Slatkin 1973, Felsenstein 1977, Kirkpatrick and Barton 1997, Behrman and Kirkpatrick 2011, Yeaman and Whitlock 2011, Le Corre and Kremer 2012). When populations are locally adapted, immigrant alleles to a given location may be poorly suited to the local environment, as these immigrants originate from populations adapted to different environments (Haldane 1956, Mayr 1963, Kirkpatrick and Barton 1997, Lenormand 2002, Polechová and Barton 2015). These alleles can impose a “migration load” on populations, reducing population size due to lower average fitness of individuals in a population (Barton 2001, Lenormand 2002, Farkas et al. 2013, Polechová and Barton 2015). Assuming organisms have a limited ability to disperse into appropriate environments (e.g. passive dispersers), migration load increases with increasing rate and spatial scale of gene flow (among other factors discussed below, Slatkin 1973, Kirkpatrick and Barton 1997, Polechová and Barton 2015).

The observation that humans are rapidly changing global environments has motivated studies of temporal changes in selection (Bay et al. 2017, Siepielski et al. 2017). Environmental change can cause population decline, extinction, or persistence via plasticity or evolution (Aitken et al. 2008). Theoretical and experimental studies have largely focused on two scenarios of environmental change: 1) a rapid, abrupt shift from a historical selection regime to a new one (Gomulkiewicz and Holt 1995, Orr and Unckless 2008) or 2) sustained change in selection through time (Pease et al. 1989, Lynch and Lande 1993, Polechová et al. 2009, Gonzalez et al. 2013). Most theoretical studies have focused on the binary outcome of whether species survive or go extinct following environmental change. For example, a number of authors have investigated factors influencing the probability of evolutionary rescue (Gomulkiewicz and Holt 1995, Orr and Unckless 2008, Bell and Gonzalez 2009, Uecker et al. 2014), which is defined as adaptation that prevents extinction following environmental change (Gonzalez et al. 2013). Still, little is known about how evolutionary response to rapid environmental change impacts abundance patterns, apart from equilibrium abundance of individual populations (Polechová et al. 2009). Despite this gap, community and ecosystem processes are strongly influenced by abundance dynamics of component species, such that understanding abundance responses to environmental change is a central goal of community and ecosystem ecology (Loreau 2010, Clark et al. 2014a). An emerging area of inquiry has investigated community evolutionary rescue, roughly defined as evolutionary rescue of multiple co-occurring species (Kovach-Orr and Fussmann 2013, Fussmann and Gonzalez 2013, Low-Décarie et al. 2015).

Among the factors that determine population response to environmental change are initial population size and genetic diversity in the trait(s) under selection. When populations are small before environmental change, they face a greater risk of stochastic extinction following environmental change (Gomulkiewicz and Holt 1995). Additionally, if genetic variants do not exist within a population that are beneficial after environmental change then a population will wait for new mutations or immigrant alleles (e.g. Orr and Unckless 2008), a scenario most relevant when adaptation is oligogenic. Alternatively, standing variation within populations may allow rapid adaptation, if adaptive variants are already present at the time of environmental change (Bonhoeffer and Nowak 1997). Such standing variation can be caused by gene flow along spatial selective gradients (Barton 2001). In particular, quantitative genetic models of local adaptation are relevant to adaptation to anthropogenic change because phenotypes involved in climate adaptation are often complex with polygenic architecture (Bay et al. 2017).

The effects of rapid environmental change on biodiversity are partly influenced by how multiple co-occurring species simultaneously respond to environment (Bradshaw 1984, Jackson and Overpeck 2000, Gilman et al. 2010, Urban et al. 2012). Typically studies of community and ecosystem responses to environmental change focus on ecological mechanisms, e.g. interspecific variation in demographic and physiological response to environment (Deutsch et al. 2008, Clark et al. 2014b, Lasky et al. 2014). For example, interspecific variation in dispersal ability is expected to have major effects on community response to environmental change, as some species are better able to track spatial shifts in environmental niches (Ackerly 2003, Gilman et al. 2010, Urban et al. 2013). However, most approaches ignore an important set of processes: intraspecific variation and evolutionary response within members of a community. Authors have studied how multiple species simultaneously evolve following environmental change using simulation (De Mazancourt et al. 2008, Vanoverbeke et al. 2015, Moran and Ormond 2015). However, many multi-species models typically focus on species that begin having niche differentiation along climate gradients (e.g. De Mazancourt et al. 2008, Price and Kirkpatrick 2009, Norberg et al. 2012, Moran and Ormond 2015), but what happens for species occupying similar climatic niches remains to be explored (Osmond and Mazancourt 2013, but see Fussmann and Gonzalez 2013).

Here I build on existing quantitative genetic theory of local adaptation (Barton 2001) and adaptation to a shifting optimum (Pease et al. 1989, Lynch and Lande 1993, Polechová et al. 2009). I reframe this theory to demonstrate the complex role interspecific variation in gene flow plays in communities due to its effect on genetic diversity, which induces migration load on populations but also causes faster adaptation (Pease et al. 1989, Polechová et al. 2009, Kremer et al. 2012). I then ask how interspecific variation in gene flow and other traits impact community dynamics following environmental change due to ecological and evolutionary processes.

## Model and Results

I start with a model of locally-adapted populations following Pease et al. (1989), Kirkpatrick and Barton (1997), Barton (2001) and Polechová et al. (2009), a stochastic version of which was studied by Polechová and Barton (2015) (referred to as the continuum of alleles model by Barton (2001)). The model I use is a deterministic model of a population with logistic growth and a quantitative trait *z* subject to hard selection with a spatially-varying selective gradient. The mean per capita reproductive rate is given by

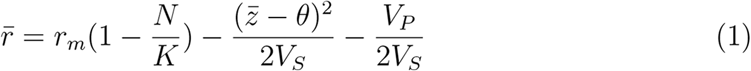

where *r*_*m*_ is population growth rate of optimal phenotype individuals at low density, *N* is census population size, *K* is carrying capacity (assumed constant through space), and *V*_*P*_ is variance of phenotype *z* (assuming a Gaussian phenotype distribution, Kirkpatrick and Barton 1997). The first term on the right-hand side of equation 1 determines a reduction in fitness due to negative density dependence. The second term gives reduction in fitness due to the mismatch between the population mean phenotype *z̄* and the local optimum *θ*, and *V*_*S*_ gives the inverse strength of stabilizing selection. Even if the population is adapted to the local optimum (i.e. *z̄* = *θ*) there still may be many maladapted individuals (i.e. *V*_*P*_ > 0), whose contribution to population mean fitness is determined by the last term in equation 1.

The optimal trait value *θ* changes in space (*x*) at rate *b* such that *θ*(*x*) = *bx* (Kirkpatrick and Barton 1997). The mean trait *z̄* at a given location *x* changes through time due to curvature of the cline in space, asymmetric gene flow (modeled as a Gaussian with standard deviation *σ*) across the cline due to spatial trends in abundance, and selection, given by the first three terms on the right hand side of equation 2, respectively (Pease et al. 1989)

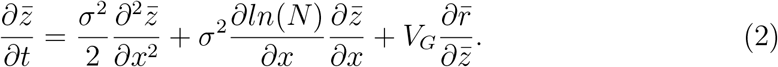

Population dynamics at *x* are given by

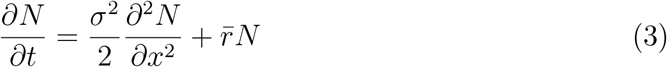

where the first term on the right-hand side gives change due to migration and spatial trends in abundance, and the second term gives change due to average individual fitness (Pease et al. 1989). Note that here there is no frequency or density-dependent selection, i.e. intraspecific competition (or apparent competition) is not dependent on *z* in any way, beyond the effects of *z* on *N*. This assumption may be well-justified for traits involved in abiotic stress-tolerance (e.g. cold or heat tolerance) where selection does not promote diversity in *z*.

Barton (2001) allowed genetic variance within a population (*V*_*G*_) to change (evolve) due to gene flow among populations. As gene flow increases, so does immigration of maladaptive genotypes into any given population. A stable equilibrium exists in this model where all populations are locally adapted along the linear environmental gradient *b*, i.e. *z̄* = *θ* at all *x* (Barton 2001). At this equilibrium, 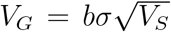. Phenotypic variance *V*_*P*_ = *V*_*G*_ + *V*_*E*_ where *V*_*E*_ is stochastic environmental variation in *z* (Barton 2001). An additional consequence of local adaptation and a linear cline in *z̄* is that 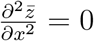 and constant population size in space, 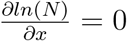. I ignore spatial boundary conditions that would result in asymmetric gene flow.

### Impacts on community structure

Two traits that ecologists commonly study are important in this model: the rate and scale of dispersal/gene flow (determined by *σ*) and reproductive rate at low density (*r*_*m*_). Maladapted immigrants depress mean fitness (known as migration load, equation 1). The equilibrium census population size (Polechová and Barton 2015) as a proportion of carrying capacity *K*, 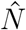, is given by

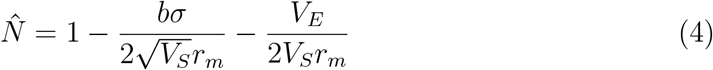

where the second term on the right gives migration load. Migration load can thus introduce uneven community structure when species differ in *σ* or *r*_*m*_. To identify the maximum *σ* capable of persistence I set 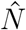 to zero and solve the inequality to obtain

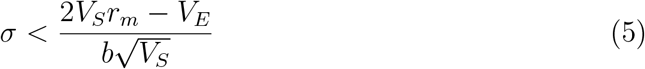

Here I am interested in complex effects of species traits that might yield unexpected results under environmental change. While greater *r*_*m*_ decreases migration load (equation 4) it does not impact the rate of adaptation (equation 2). However, gene flow, *σ*, plays a more complex role.

To study how interspecific variation in *σ* could structure communities along spatiotemporal environmental gradients, I now consider a community of species that vary only in *σ* (but not other parameters e.g. *K*, *V*_*S*_, *V*_*E*_). For mathematical convenience I start with communities lacking species interactions. I follow with simulations that introduce competition among species.

In the Barton (2001) model, greater *σ* increases *V*_*G*_ and migration load and thus decreases equilibrium population size. From equation 4, the proportional reduction in 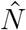 due to migration load is equal to 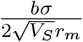. I differentiate with respect to *σ* to obtain

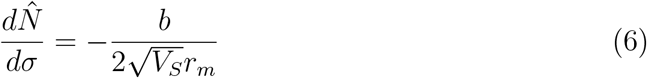

which gives the slope of species equlibrium abundance versus gene flow. Thus the species abundance distribution for a community (McGill et al. 2007) could be obtained using the distribution of *σ* and applying equation 6. The parameters on the right of equation 6 are each constrained to be positive so that when holding these constant across species of varying *σ* there is a negative relationship between *σ* and 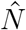. The effect of migration load is stronger and the abundance distribution is steeper as the selective gradient *b* is steeper.

### Abrupt environmental change and transient community turnover

The interesting effects of gene flow in a community context arise from the dual role of *σ* following environmental change. Greater *σ* can have a fitness benefit when population mean traits differ from the optimum, *z̄* ≠ *θ*, such as in populations that have experienced recent environmental change (Polechová et al. 2009, Kremer et al. 2012) or populations colonizing new environments. Greater *σ* proportionally increases 201 *V*_*G*_, which proportionally increases the speed of adaptation (third term on right-hand side of equation 2). I studied the effect of *σ* on population and community dynamics using numerical simulations. I simulated populations with non-overlapping generations governed by discretized versions the above equations. Simulations were intialized with locally-adapted populations at equillibrium population size, 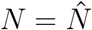 and *z̄* = *θ*.

I chose biologically plausible parameter values (although below I study other values): *b* = 0.05, *V*_*S*_ = 1, *V*_*E*_ = 0.05, *r*_*m*_ = 0.5, *x* = 0 and thus *θ* = 0 (Polechová and Barton 2015). I then imposed an instantaneous change in *θ* such that a new phenotype, *θ*\* = 1, was optimal, and the change in selection was the same at all locations, i.e. the slope *b* of the spatial gradient did not change, *θ*\*(*x*) = *bx* + 1 (Figure 1). This scenario is mathematically convenient because all populations experience the same relative change and dynamics and thus no spatial trend in abundance emerges nor does the cline in *z̄* change. If a system begins at locally-adapted equilibrium, a change in *θ* by the same amount at all locations *x* will leave *V*_*G*_ unchanged because the slope of the cline in *z̄* is unchanged (see equation 10B in Barton 2001).

**Figure 1.**
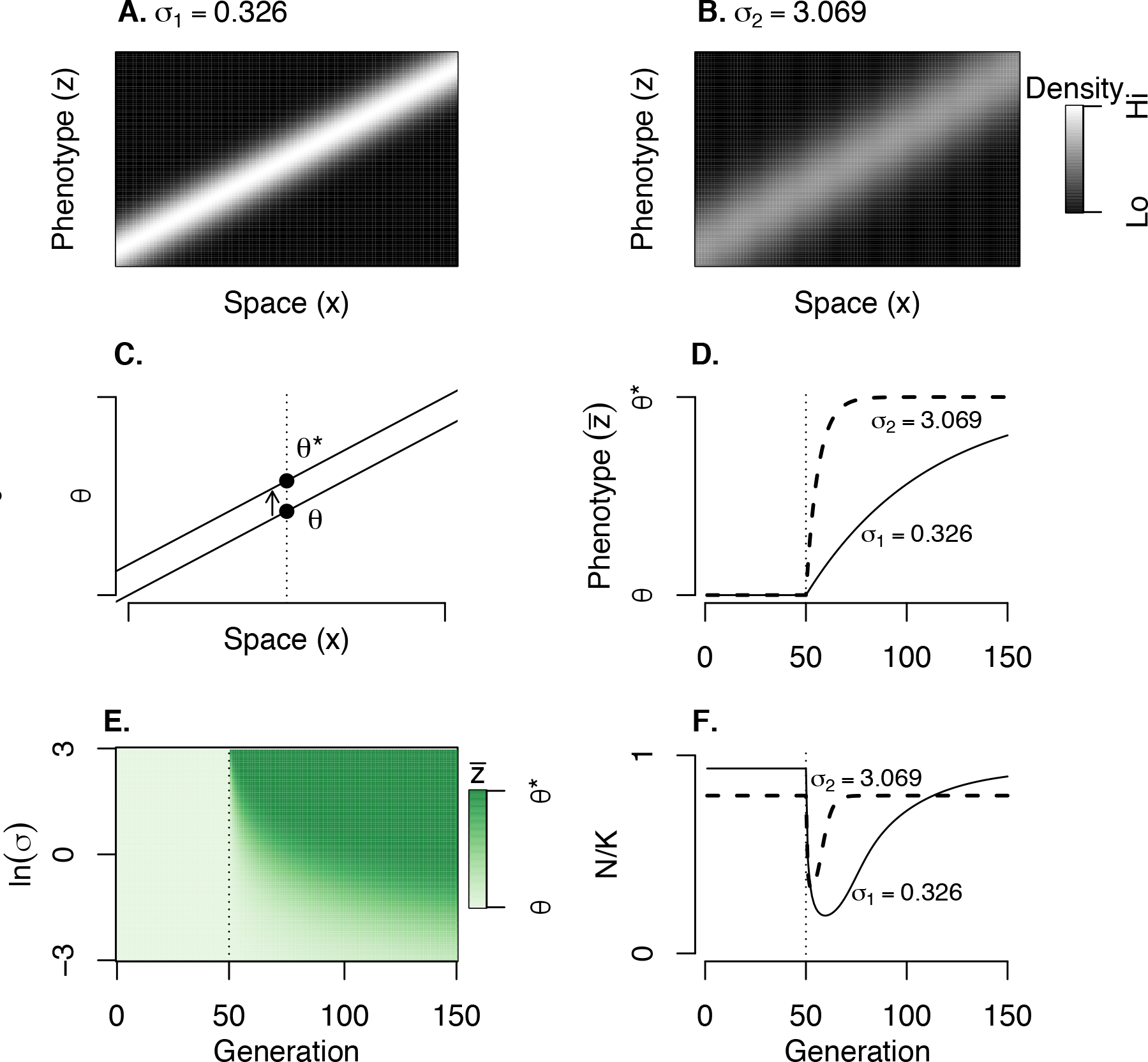
In a locally adapted system, interspecific variation in *σ*, determining the rate and scale of gene flow, determines differences in *V*_*G*_ and rate of adaptation. Here there is no interspecific competition. Species with low (A) and high (B) *σ* are subject to the same selective gradient *b* (favoring an increase in phenotype value through space from left to right) and all populations are locally adapted. (B) The high *σ* species has higher diversity of the trait under selection within populations (*V*_*G*_) (evident via a thicker gray smear for any given location along the x-axis) due to maladaptive immigration. (C) An instantaneous change in optimal phenotype from *θ* to *θ*\* occurs at generation 50. (D) The higher *σ* species adapts to the new optimum faster (D), and (E) when comparing 100 species with a range of *σ* values. (E) White is the optimal trait prior to the change, green is the optimal trait following the change. (F) Faster adaptation by a high *σ* species compared to a low *σ* species leads to transient community turnover. Parameter values (unless otherwise noted) were *b* = 0.05, *V*_*S*_ = 1, *V*_*E*_ = 0.05, *r*_*m*_ = 0.5, and *θ*\* − *θ* = 1.

I first compare evolution of *z* for two non-interacting species differing only in *σ*(*σ*_1_ = 0.326 and *σ*_2_ = 3.069). Both species were subject to the same selective gradient *b* = 0.05 and the clines in the mean phenotype *z̄* of the two species were equal before environmental change, but with the second species having greater variance within any local population (i.e. greater *V*_*G*_, Figure 1). The high gene flow species rapidly adapts to *θ*\* with the low *σ* species lagging far behind (Figure 1D).

Faster adaptation following a shift in environment will lead to more rapid recovery of population mean fitness. Although species with high *σ* are less abundant than low *σ* species in communities in a stable environment (eqn. 4), the faster adaptation of high *σ* species can allow them to increase their relative abundance following an environmental change. These two example species (*σ* = 0.326 and *σ* = 3.069, respectively) exhibit a transient reversal in relative abundance as the high *σ* species is more abundant for an interval following the environmental change (Figure 1F). The reversal is transient because the stable environment after change again favors low *σ*. This transient shift to species with high *σ* and back to species with low *σ* also emerges if this system is subjected to ecological disturbance (Figure S4). Thus the predicted patterns of eco-evolutionary turnover from this model may follow patterns of ecological succession, 233 albeit due to different mechanisms.

I now introduce species interactions into the model. In a diverse community of interacting species that vary in gene flow one can ask how composition might shift due to different evolutionary responses. Species interactions could change the relative importance of some of the processes studied previously. For example, interspecific competition could depress the mean fitness of species, pushing them closer to extinction, and also exacerbate relative population differences. Here I build on the previous quantitative genetic models to simulate species within a community competing against each other, using the Lotka-Volterra form. Instead of equation 1, I used a discrete time version of the following

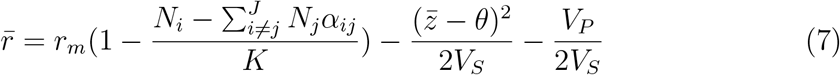

where *N*_*i*_ is the population size of the focal species *i* and there are *J* total competitor species each with population sizes of *N*_*j*_. *α*_*ij*_ determines the strength of interspecific competition. Interactions were symmetric among species such that all *α*_*ij*_ = *α*_*ji*_. Note that per equation 2, I assume adaptation is not influenced by such competitive interactions (i.e. competition does not influence 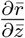; *α*_*ij*_ is unrelated to *z*_*i*_and *z*_*j*_, cf. Osmond and Mazancourt 2013, Fussmann and Gonzalez 2013).

I simulated communities with a log uniform distribution of *σ* values across 100 species under the same conditions as the two previous species, but now with interspecific competition (*α*_*ij*_ = 0.1). I initiated species at a low abundance (*N* = 10^−5^), but then allowed 500 generations for population growth with interspecific competition and constant *θ*, before imposing change in *θ* and simulating for 500 more generations. I calculated which species was most abundant at each time point.

Under equilibrium, the species with lowest *σ* has highest *N* (Figure 2). Equation 4 gives 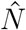 when there is no interspecific competition. In a diverse community, all species experience approximately equal effects of interspecific competition and thus the relative differences among species in 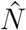 remain approximately the same, albeit with a decrease in the maximum *σ* capable of persisting (Figure S2). Following an instantaneous shift to *θ*\*, higher *σ* species dominate but gradually give way to lower *σ* species. However, the time required for poor dispersers to adapt can be long given their slow rate of adaptation (Figure 1E). This interspecific variation in adaptation following environmental change will likely have impacts on the distribution of traits in a community, which is often of interest to community and ecosystem ecologists (Muscarella and Uriarte 2016, Šímová et al. 2018). For example, ecosystem function may be influenced by the mass-averaged functional traits in a community (Grime 1998). In the Online Supplement I show how interspecific competition causes community 8 mean *z* to more quickly approach *θ*\* as fast-adapting high *σ* species supress the initially abundant low *σ* species, especially under a scenario of abrupt environmental change (Figure S1).

**Figure 2.**
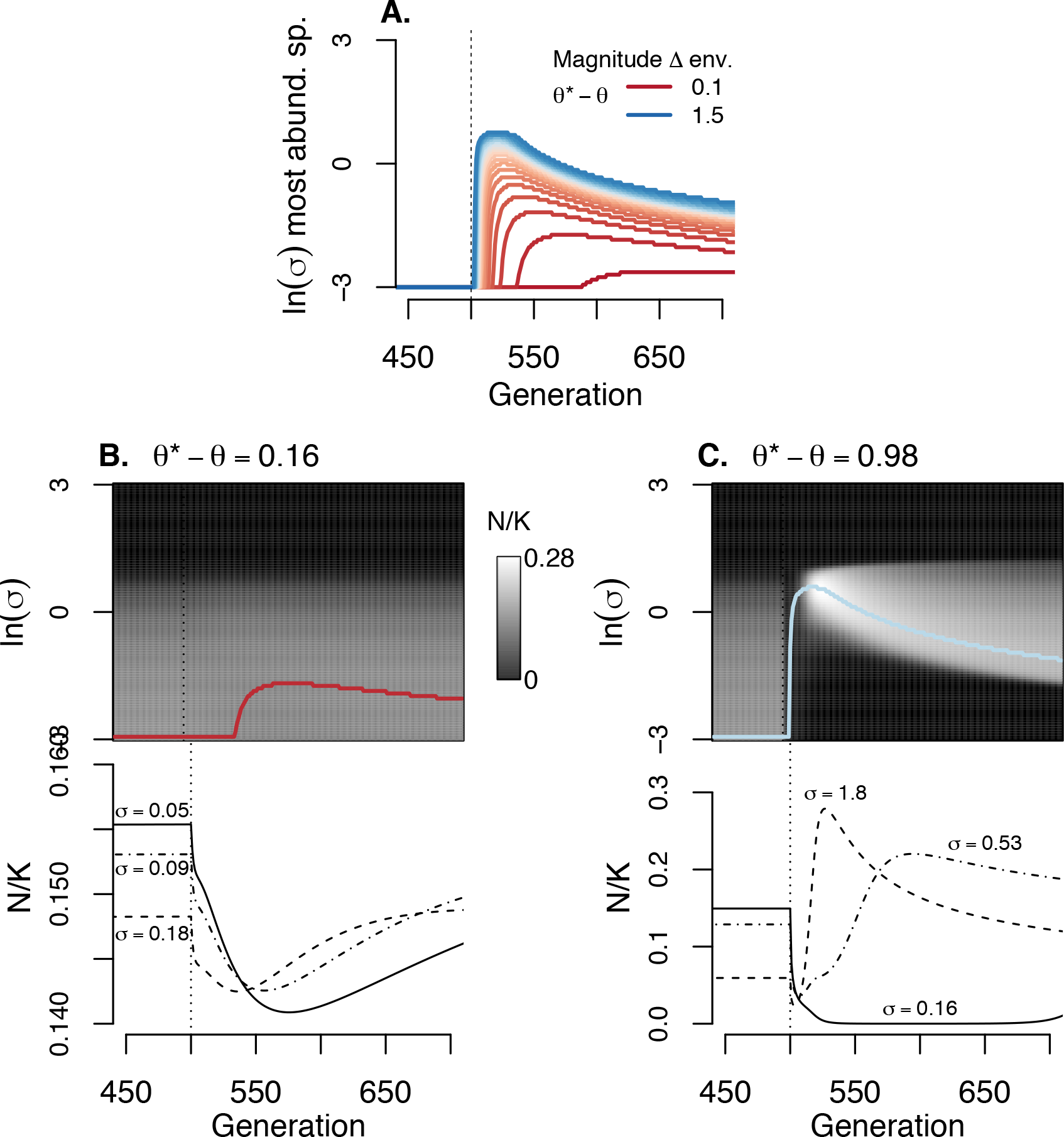
How the magnitude of environmental shift affects the magnitude of community turnover in the presence of interspecific competition among *J* = 100 species (all *α*_*ij*_ = 0.1). (A) In general, the greater the environmental change, the higher the *σ* of the most abundant species soon after the environmental change. (B-C) Example community and individual species trajectories for specific magnitudes of environmental change. (B) When environmental change is smaller, a lag between environmental change and change in species rank abundances can occurr. (C) Under larger environmental changes, high *σ* species can exhibit population spikes as they are released from competition with low *σ* species. Populations are at equilibrium and adapted to *θ* for the first 500 generations, when an instantaneous environmental change to *θ*\* occurs. Parameter values (unless otherwise noted) were *b* = 0.05, *V*_*S*_ = 1, *V*_*E*_ = 0.05, and *r*_*m*_ = 0.5.

Because the transient advantage of higher *σ* species comes from their faster approach of *z̄* to *θ*\* (Equation 2), the magnitude of environmental change might influence the degree of community turnover. Under a weak shift in *θ*, the benefit to adapting faster for high *σ* species is low (Figure 2). When the magnitude of the environmental shift is large, community turnover (as defined as which species dominate following the environmental shift) is also large. Notably, subtle shifts in environment lead to subtle, though delayed changes in the most dominant species (blue lines in Figure 2A). This lag emerges because when a species starts with greater *N* in a constant environment the differences between species in maladaptation take time to erode the initial advantage (Figure 2). Despite the lag in reversal of species rank abundances, the differences among species in *r* are quickly evident in the form of differences in 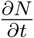 (i.e. there is rapid emergence of differences among species in slope of *N* trajectories, Figure 2B).

#### The strength of species interactions

To evaluate how the strength of species interactions can change eco-evolutionary response to environmental change, I simulated communities with different values of *α*_*ij*_ and compared results. Comparing scenarios with *J* = 100 species and *α*_*ij*_ equal to 0, 0.01, or 0.1, showed little effect on turnover in the most abundant community member (Figures 2 and S3). Thus the main effect of adding weak to modest pairwise interspecific competition in a diverse community was to reduce the maximal *σ* capable of persisting. Concordantly, variation in the magnitude of abrupt environmental change had similar impact on community dynamics, as measured as *σ* of the most dominant species, across these values of *α*_*ij*_. Note that although e.g. *α*_*ij*_ = 0.01 means individual species interact weakly, the presence of many other species (e.g. *J* = 100) in the community can result in substantial competition in aggregate.

I also simulated ten strongly competing species (*α*_*ij*_ = 0.75) and found substantial differences in community dynamics. Here, competition had a stronger effect on how the *σ* of the most abundant species changed with time (Figure S3). Competition resulted in dominance of species with relatively lower *σ* shortly after environmental change. There were even stronger effects of competition on the dynamics of individual species. In the presence of this strong interspecific competition, low *σ* species that have relatively lower abundance following environmental change remained supressed for longer periods of time and at very low densities (Figure S3). Close inspection of the results showed that these low *σ* species that reached low density following environmental change were on an upward population trend at the end of simulations. Thus the dominance of higher *σ* species was still transient, though with a much slower return to the pre-environmental change equillibrium 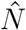. Note that my deterministic simulations lack stochastic extinction, which is likely a major problem for populations at very low density. Higher *σ* species can actually see increased absolute abundance following environmental change, despite going from being locally-adapted to being maladapted (Figures 2C and S3). This surprising change results from the release from competitive supression by low *σ* species. This spike is particularly pronounced for intermediate to high *σ* species that have a good balance of adaptability versus migration load (Figure S5).

#### Modifiers of the tradeoff between migration load versus adaptability

I next studied how factors that mediate the tradeoffs associated with *σ* (migration load versus speed of adaptation) impact community dynamics. Migration load is ameliorated under shallower environmental gradients (lower *b*), though low *b* also reduces *V*_*G*_ and hence the rate of adaptation. In nature, the slope of environmental gradients varies in space and is thought to be an important driver of biodiversity patterns (Yeaman and Jarvis 2006). Under low *b*, there will be predominantly gene flow between like environments. The slope of the curve relating species abundance to gene flow 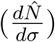 is proportional to *b* thus lower *b* will result in a shallower abundance curve, i.e. a more even community. That is, migration load is reduced and species differing in *σ* have similar abundances at equilibrium.

When I varied *b*, the most obvious impact is on the magnitude of community turnover following environmental change (Figure 3A, simulations with same parameter values as above except *J* = 100 species and all *α*_*ij*_ = 0.1). Immediately after the environmental change, high *σ* species dominate when *b* is low. Note that when *b* is low, differences in abundance of species differing in *σ* are subtle due to low migration load, though there is relatively high turnover in which species are most abundant following the environmental change. When *b* is high, the environmental change results in turnover favoring species of intermediate *σ*. Surprisingly, the temporal change in relative species abundances following the environmental change happens at a similar rate regardless of *b* (lines in Figure 3A have similar trajectories following environmental change), although higher *b* results in faster return to equilibrium because the initial community turnover was less. The consistency of the rate of community turnover is likely due to species proportional differences in *V*_*G*_ and rate of adaptation being constant despite differences in *b* (equation 2).

**Figure 3.**
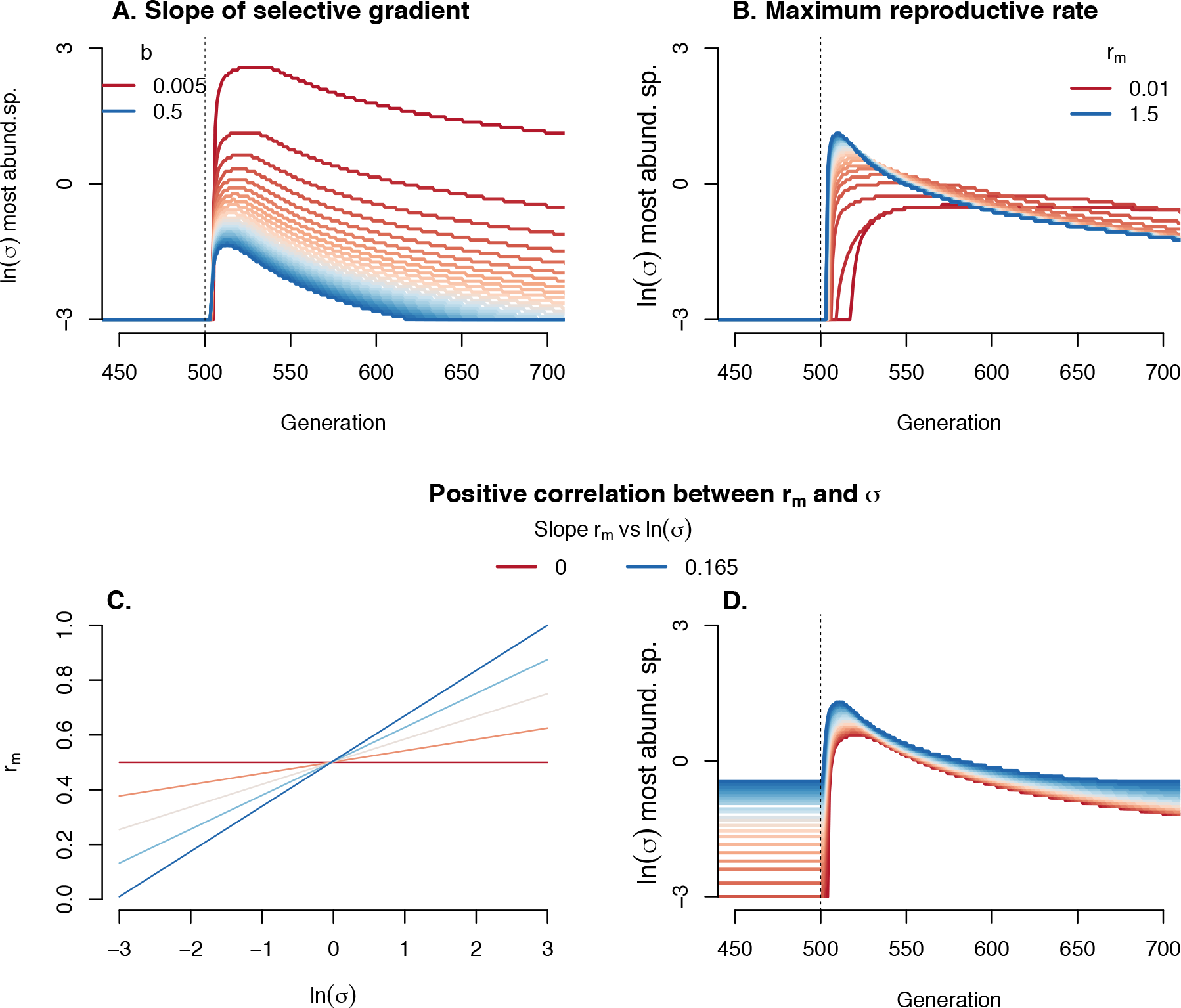
Modifiers of the tradeoff between migration load versus adaptability in the presence of interspecific competition among *J* = 100 species (all *α*_*ij*_ = 0.1). (A) The slope of the selective gradient (*b*) affects tradeoffs associated with *σ* and community turnover following an abrupt environmental change. Greater *b* results in dominance by intermediate *σ* species folowing abrupt environmental change (imposed after 500 generations). Lower *b* allows higher *σ* species to briefly dominate, although in these scenarios migration load is low and abundance at equillibrium 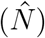 under stable environments is only weakly related to *σ*. (B) Greater reproductive rate at low density *r*_*m*_ ameliorates migration load and affects community turnover following an abrupt environmental change. Greater *r*_*m*_ results in an initially greater commuity turnover because lower migration load allows high *σ* species to leverage their faster adaptation. Lower *r*_*m*_ increases migration load, limits the ability of high *σ* to take advantage of their faster adaptation, but also slows the rebound of eventually dominant low *σ* species. (C-D) Correlation between *r*_*m*_ and *σ* affects community turnover following an abrupt environmental change. Greater correlation results in dominance by intermediate (as opposed to low) *σ* species at equillibrium under constant environments, and hence less turnover following environmental change. Parameter values (unless otherwise noted) were *b* = 0.05, *V*_*S*_ = 1, *V*_*E*_ = 0.05, and *θ*\* − *θ* = 1, and *r*_*m*_ = 0.5.

Barton (2001) and Polechová and Barton (2015) investigated how faster change in environments at range margins, i.e. increasing magnitude of *b*, impacts local adaptation. My results on how *b* influences community turnover due to differential evolutionary response to environmental change may apply to such changes in *b* in space. The present model can be applied assuming that the rate of change in *b* is subtle, such that ∂ *z̄*/∂*x* remains approximately linear. If *b* is sharper at range margins (for an assemblage of species, this would correspond to ecotones at the margin of ecoregions, for example along very steep altitudinal gradients), migration load would be stronger at margins and would have a stronger influence on community composition at equilibrium (i.e. steeper 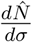). However, following environmental change, the change in species rank abundance will be greater in the range core (low *b*) while there would be lesser change in species rank at range margins (high *b*).

Migration load is also ameliorated by high *r*_*m*_ (equation 4), thus *r*_*m*_ may impact eco-evolutionary community dynamics. Greater *r*_*m*_ reduces the effects of maladaptive immigration on 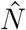 and allows for persistence (i.e. 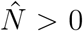) of species with higher *σ* (inequality 5). My simulations showed opposing effects of *r*_*m*_ on community dynamics. When *r*_*m*_ is low, high *σ* species cannot persist and thus the magnitude of community turnover is lower. However, because *r*_*m*_ is low, the recovery of species from low density is slow, and the community is dominated by relatively higher *σ* species for a long period of time (Figure 3B). By contrast, high *r*_*m*_ allows for the existence of high *σ* species and the rapid environmental change causes strong, but shorter lived, community turnover.

Interspecific trait variation is often correlated across multiple trait axes, corresponding to ecological strategies and life histories. Thus it is unlikely that natural variation in *σ* is independent of other traits. To study the impacts of trait covariation, I simulated the situation where *σ* and *r*_*m*_ positively covary such that higher gene flow species also exhibit higher per capita population growth when rare. For example, plants with high reproductive rates tend to have greater dispersal distances (Beckman et al. 2018). I simulated a positive log-linear relationship similar to the empirical relationship for 141 species observed by Beckman et al. (2018), *r*_*m*_ = *a* + *cln*(*σ*), where *a* is an intercept and *c* determines the rate at which *r*_*m*_ increases for species of higher *ln*(*σ*). This correlation has opposing effects on migration load: *r*_*m*_ decreases load but *σ* increases load (equation 4). Thus intermediate *σ* species have greatest abundance at equillibrium (Figure 3D). Notably, this correlation between *r*_*m*_ and *σ* leads to weaker eco-evolutionary community turnover because intermediate *σ* species were already dominant before environmental change so their dominance shortly after environmental change means the community is relatively consistent.

### Community turnover under sustained environmental change

Temporal environmental change can take any functional form. In the previous section I simulated an instantaneous shift in environment that then stabilized (Gomulkiewicz and Holt 1995, Orr and Unckless 2008). Alternatively, environments may undergo more gradual sustained directional shifts. This scenario has been analyzed previously by Pease et al. (1989), Lynch and Lande (1993), and Polechová et al. (2009). Here, I build on this framework by explicitly considering the role of gene flow on population dynamics in this scenario. In the Lynch and Lande (1993) single-species model, the optimum *θ* changes at a rate *k* per unit time *t*, so that *θ*\*(*x*, *t*) = *bx* + *kt* (Polechová et al. 2009). After a enough time has passed to allow for a balance between adaptation versus the shifting optimum, the mean trait (*z̄*) at location *x* lags behind the optimum a stable distance, which Lynch and Lande (1993) calculated as equal to 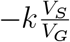. In the present model, greater *σ* increases *V*_*G*_ and causes lower lag in *z̄* behind the shifting optimum. Substituting the Barton (2001) equation for *V*_*G*_ in a system at locally-adapted equilibrium into the previous expression results in a lag in *z̄* equal to

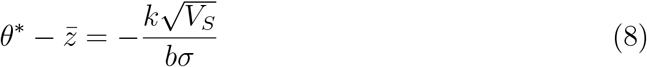

i.e. lag in *z̄* for a given species is proportional to *σ*^−1^ (Polechová et al. 2009 identified this expression in a population genetic model of this scenario). Thus stronger stabilizing selection reduces the lag, though to a lesser degree than identified by Lynch and Lande (1993, 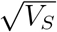 versus *V*_*S*_, Kremer et al. 2012). This is because when stabilizing selection is stronger (low *V*_*S*_) the fitness advantage of adapted genotypes is higher but stronger stabilizing selection also reduces *V*_*G*_ from immigration, slowing adaptation.

Lynch and Lande (1993) also derived the critical rate of environmental change above which populations go extinct (ignoring stochasticity) as 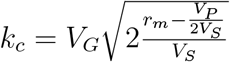 (see also Polechová et al. 2009). I substitute the Barton (2001) equation for *V*_*G*_ in a locally adapted system into the previous equation to obtain

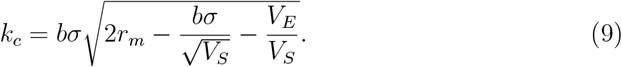

This equation shows how *k*_*c*_ is non-monotonically related to *σ*, i.e. *k*_*c*_ is greatest for intermediate values of *σ* (Polechová et al. 2009). To determine how the shifting optimum impacts community structure as *t* becomes large, I substituted the lag in *z̄* to the previous equation for 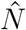 (equation 4), while still using the single-species model of >Lynch and Lande (1993) (i.e. *α*_*ij*_ = 0). Thus at equilibrium trait lag under a shifting environment

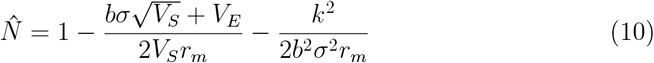

where the first substracted term includes migration load, which worsens with *σ*, while the second substracted term gives the lag load, which is ameliorated by *σ*. These opposing effects result in species with intermediate values of *σ* and hence *V*_*G*_ being most abundant (Figure 4, Polechová et al. 2009). Differentiating with respect to *σ* gives

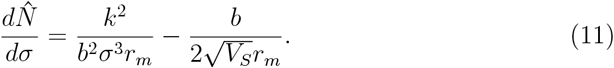

The maximum 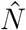 is attained by species with *σ* that cause the right hand side of equation 10 to equal zero, i.e. the *σ* with maximum 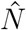 is equal to 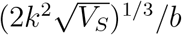. Note this expression equals zero when *k* is zero, thus consistent with results on locally adapted systems in constant environments where *σ* = 0 is favored due to lack of migration load (equation 6). Thus greater rates of environmental change through time (*k*) favor higher *σ* species, but at a decreasing rate 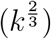.

**Figure 4.**
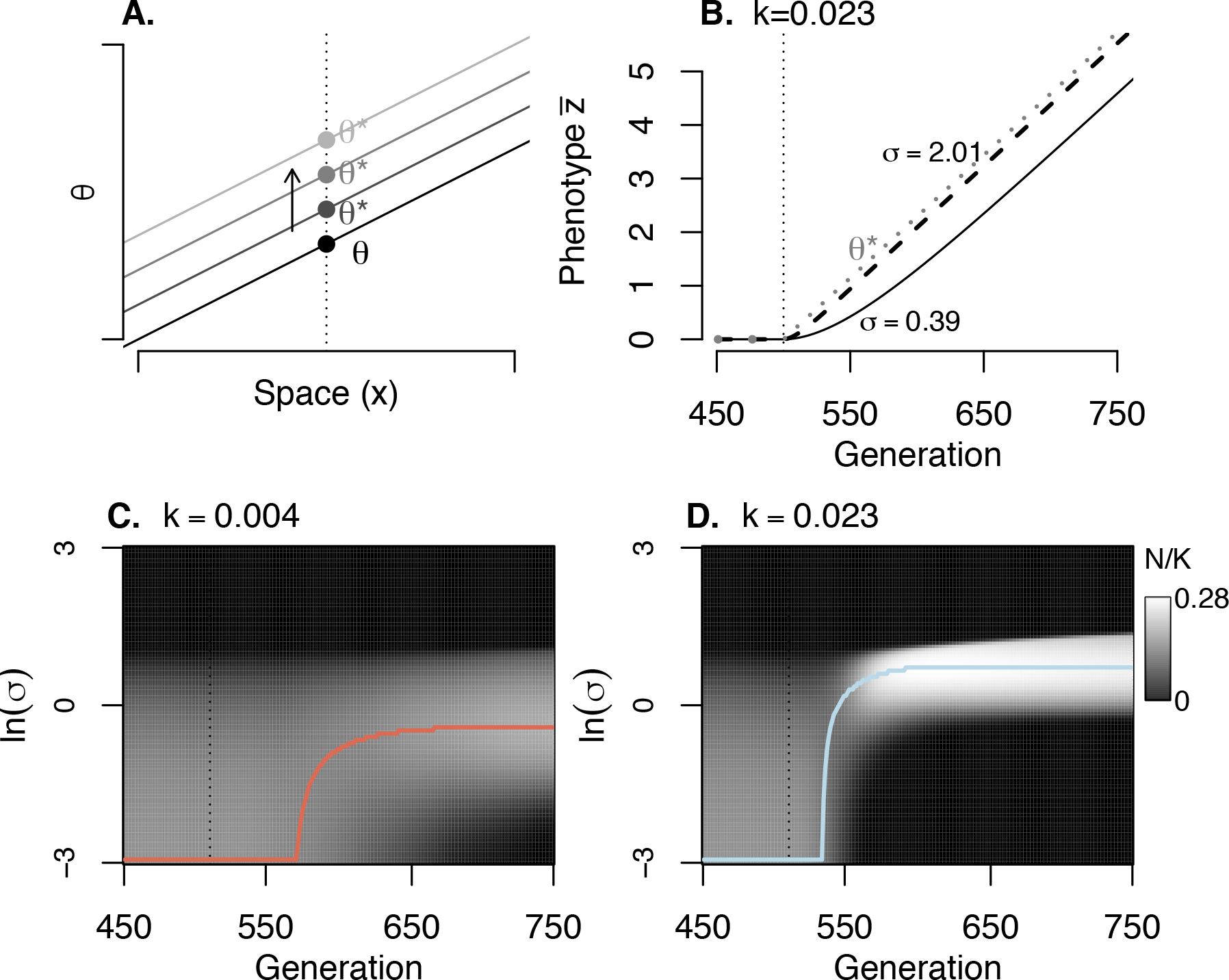
Effects of a sustained environmental change (i.e. a change in *θ* through time), in the presence of interspecific competition among *J* = 100 species (all *α*_*ij*_ = 0.1). (A) Illustration of the scenario of shifting *θ* across all locations, from a historical *θ* to which species were locally adapted, to *θ*\*. (B-D) Environment is constant (constant *θ*) until the vertical dashed line at which point *θ* changes at a constant rate *k*. (B) Illustration with *k* = 0.023 of evolution of *z̄* for two example species differing in *σ*, compared to the shifting optimum (*θ*). (C-D) Population size trajectories for all 100 species under different rates of change, colored lines correspond to the colors for a given *k* in Figure 5. Parameter values (unless otherwise noted) were *b* = 0.05, *V*_*S*_ = 1, *V*_*E*_ = 0.05, and *r*_*m*_ = 0.5.

In this scenario of sustained environmental change, steepening selective gradients (higher *b*) results in a lower *σ* having maximum 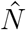 (equation 1), more similar to the situation in a constant environment. Thus these results are similar to those following an abrupt change in environment: at range margins or ecotones where *b* may be steeper, the magnitude of change in the most abundant species will be less, compared to where *b* is shallower.

#### The strength of species interactions

I also simulated how interspecific competition impacts the community response to a sustained environmental change. I used the same model of species interactions as described above (equation 7) under the scenario of shifting *θ* at rate *k* throught time. I simulated diverse communities of species (*J* = 100) with different values of *α*_*ij*_: 0, 0.01, and 0.1. I found that the *σ* of the dominant species under environmental change was largely the same regardless of these levels of interspecific competition (Figure 5). When increasing interaction strength (*α*_*ij*_ = 0.75) in less diverse communities (*J* = 10), I found similar patterns comparing *α*_*ij*_ = 0.75 to *α*_*ij*_ = 0 in terms of which species were most abundant through time (Figure 5). However, this similarity obscured substantial effects of competition on the trajectories of individual species. Under interspecific competition, the most abundant species had greater relative abundance advantages. Interestingly, in scenarios with interspecific competition, higher gene flow species often showed dramatic increases in absolute abundance following the initiation of environmental change (Figures 4 & 5). In these simulations, low *σ* species were supressed by environmental change and this allowed increased abundance of higher gene flow species better able to adapt to shifting environments. The increases in abundance by higher *σ* species were often short lived or delayed well beyond the initiation of environmental change, associated with the slow decline of intermediate (but less than optimal) *σ* species (Figures 4D & S6). The timing of these population spikes had a non-linear relationship with *σ* (Figure S6), such that low *σ* species were most abundant before environmental change began, while higher *σ* species exhibited spikes in abundance in sequence, until the highest *σ* species that were unable to persist because of high migration load (see examples in Figures 4D & 5C). Fast rates of environmental change also had the effect of reducing the diversity in *σ* in the community (compare Figure 4C and 4D). Under slow change (low *k*) a wider range of *σ* persisted. Under faster change, the loss of low *σ* species was not balanced by increases in the highest *σ* species, which were still limited by migration load.

**Figure 5.**
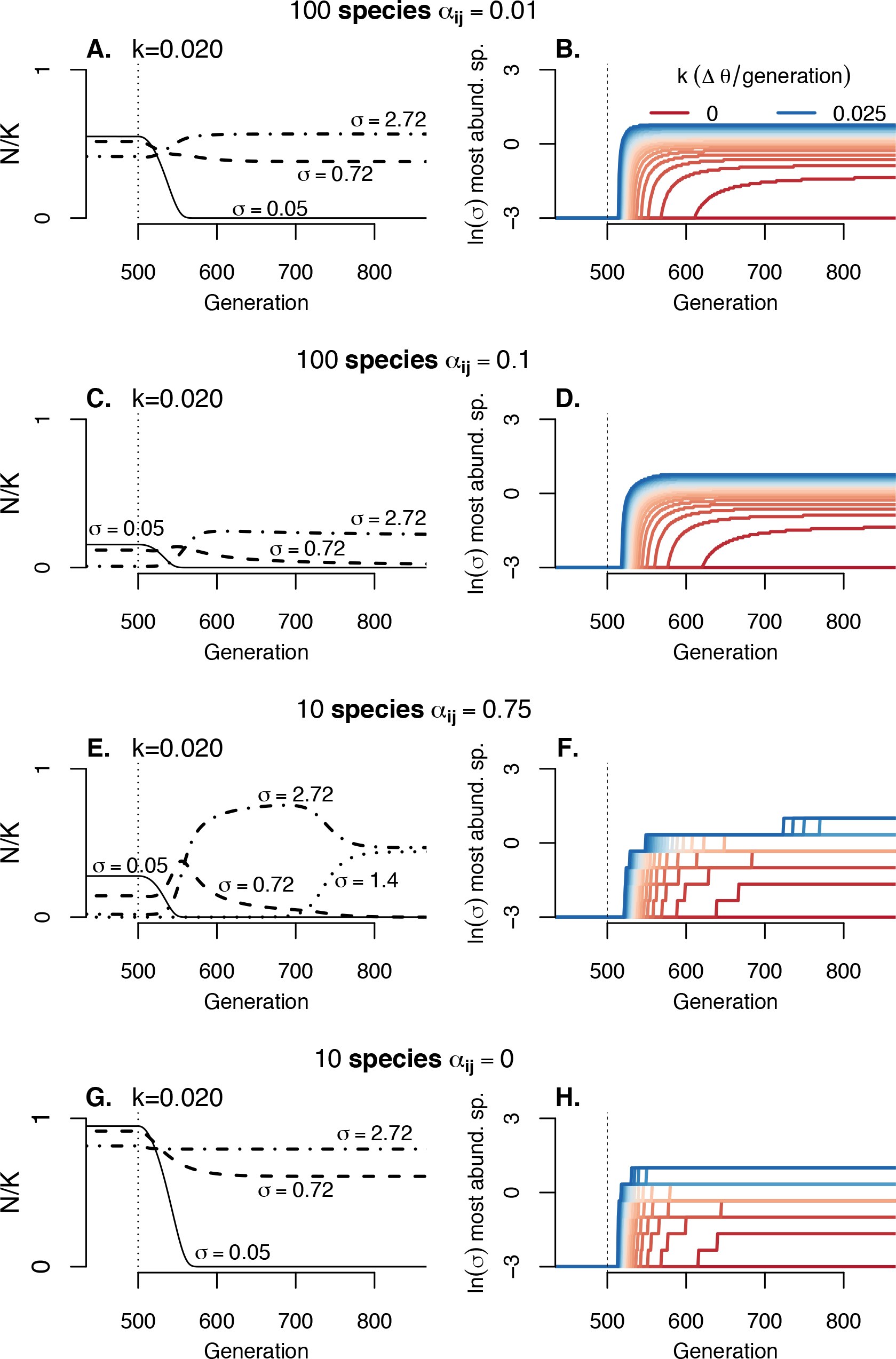
Effects of a sustained environmental change and variation in interspecific competition, with example species highlighted in each scenario. Left panels show trajectories of 79 individual example species while right panels show which species dominate communities through time. (A-D) Diverse communities with variable interspecific competition, having same species as the communities shown in Figure 4. (E-F) A community with fewer species and strong interspecific competition and (G-H) a community composed of the same species as (E-F) but with no interspecific competition. Vertical dashed line indicates beginning of environmental change at generation 500. Parameter values (unless otherwise noted) were *b* = 0.05, *V*_*S*_ = 1, *V*_*E*_ = 0.05, and *r*_*m*_ = 0.5.

### Ecosystem resilience and interspecific interactions

The increased absolute abundance exhibited by many intermediate to high *σ* species under interspecific competition and environmental change may have important community and ecosystem-level consequences. For example, biodiversity can impact ecosystem function when species exhibit compensatory population dynamics through time, stabilizing ecosystem-level processes (Micheli et al. 1999, Loreau 2010).

I quantified biomass resilience using approaches specific to each scenario of environmental change. For abrupt change, I calculated the time (number of generations) until the community regained 75% of the biomass seen at equilibrium before the environmental change. For sustained change, I calculated the biomass in the final generation of simulations (500 generations following the initiation of change - when populations had stabilized) as a proportion of the biomass under stable environments.

In both cases, simulations showed that communities with stronger interspecific com-465 petition also showed greater resilience under strong environmental change and mal-adaptation. In diverse communities with weak interspecific competition, biomass either returned faster or was maintained at higher relative levels, compared to similar communities without interspecific competition (Figure 6). Communities with fewer species (10 species) but stronger interspecific competition exhibited even greater resilience relative to comparable communities without interspecific competition, under both scenarios of environmental change. This resilience is clearly due to increases in abundance of high *σ* species, which were released from competitive supression by previously dominant but slow adapting low *σ* species, and which themselves adapt to changing environments rapidly (Figures 5 & 6).

**Figure 6.**
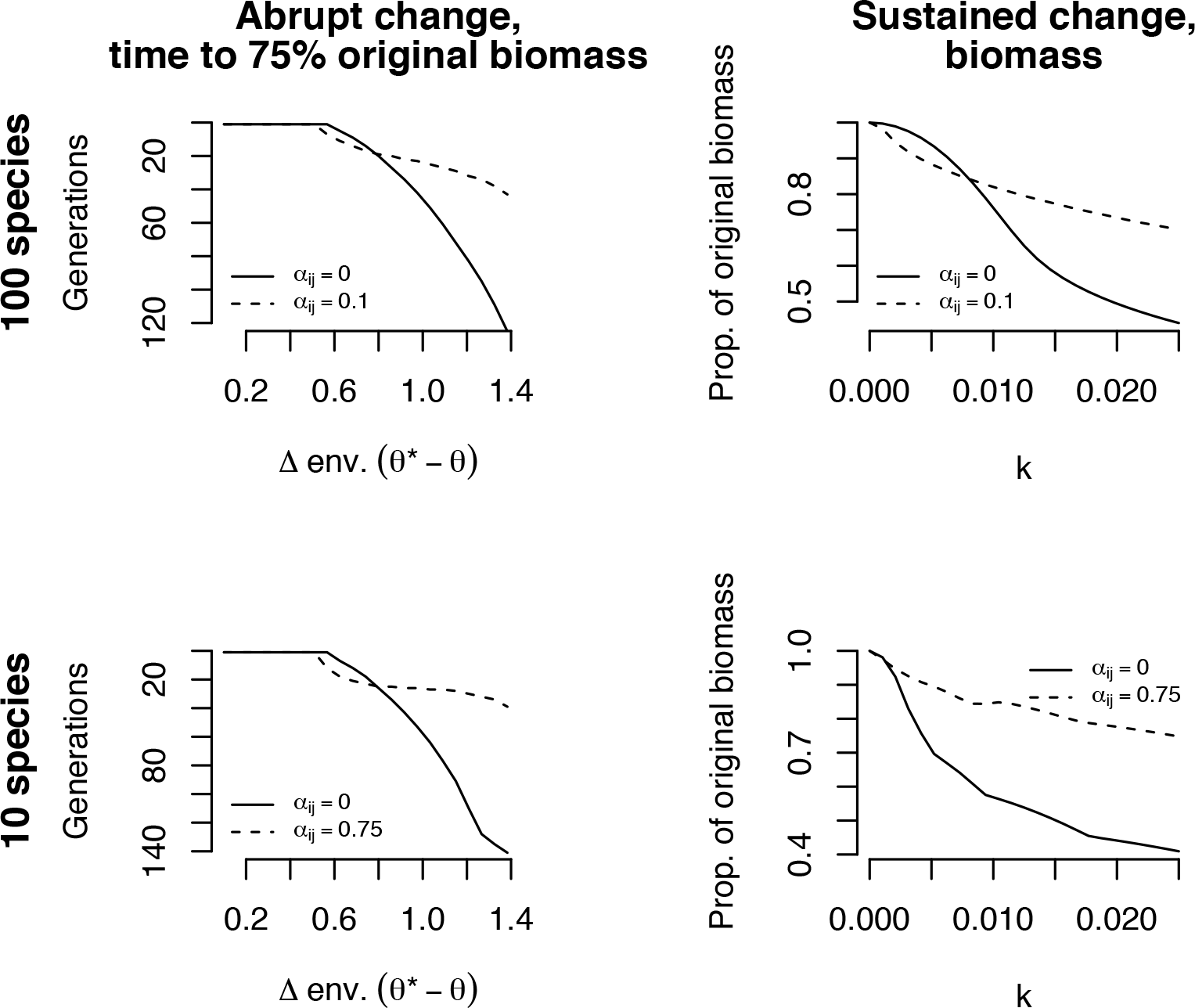
Communities with interspecific competition are more resilient to environmental change, measured in terms of (left panels) time to return to 0.75 of pre-environmental change biomass or (right panels) biomass in 500th generation under sustained linear temporal change. Note that in left panels the y-axis is reversed for comparability with right panels. Biomass is measured as the total number of individuals of all species. For left panels, *σ*\* − *σ* = Parameter values (unless otherwise noted) were *b* = 0.05, *V*_*S*_ = 1, *V*_*E*_ = 0.05, and *r*_*m*_ = 0.5.

## Discussion

Evolutionary genetic theory is a rich source of hypotheses for how life history affects evolution. On this rapidly changing planet, understanding and predicting evolutionary responses environmental change will be particularly valuable (Gienapp et al. 2017, Bay et al. 2017). Molecular data are providing a deeper view of the differences among species in population genomic patterns (e.g. Romiguier et al. 2014). The present is ripe for studying how interspecific trait differences impact evolutionary response to environmental change and the consequences for communities and ecosystems. Here, I took existing quantitative genetic models of adaptation (Lynch and Lande 1993, Barton 2001, Polechová et al. 2009) and showed how interspecific trait variation gives rise to differences in genetic diversity with non-monotonic effects on community structure and dynamics. Many previous studies of evolutionary rescue have largely focused on thresholds beyond which populations go extinct under environmental change (Lynch and Lande 1993, Gomulkiewicz and Holt 1995, Bell and Gonzalez 2009, Uecker et al. 2014). Even if populations of most species in a community are able to avoid extinction under environmental change, my results highlight how communities may change drastically in composition and function. Dominant species can become rare and rare species can become dominant (Figure 5). This turnover has important consequences for community diversity and ecosystem function.

In general, eco-evolutionary community inversions (i.e. reversals in relative abundances) may arise in any system where there is a negative or complex relationship between 6 population size and adaptability to environmental change. In my model, these changes are driven by the fact that initially numerically abundant species are more maladapted for longer periods of time following environmental change. Genetic variance has a major influence on the rate of adaptation, but other traits, such as generation time, vary among species in communities and may also result in eco-evolutionary community turnover. For example, parasites may have shorter generation time than hosts, allowing parasites to adapt faster to abiotic environmental change. Both vertebrate hosts (Fraser 2013) and their parasites (Sternberg and Thomas 2014) can be locally-adapted along temperature gradients, though parasites might adapt to climate change faster than hosts. Alternatively, when census population size is positively related to genetic variance in a trait under selection (Frankham 1996), evolutionary responses to environmental change may reinforce the ecological responses, reducing community diversity.

I identified a transient benefit to high gene flow following an abrupt environmental change, due to faster adaptation. In their experimental microcosm study, Low-Décarie et al. (2015) demonstrated how gene flow was key to the eco-evolutionary recovery of soil microbial communities responding to a novel herbicide. Studies of genetic variation (Lande and Shannon 1996) from dispersal (Polechová et al. 2009, Blanquart and Gandon 2011) or mutation (Taddei et al. 1997) have yielded similar results. When environment is constant, low mutation rates are favored, though mutator lineages have transient benefits when they find adaptive mutations (Taddei et al. 1997). Additionally, fluctuating environments can favor higher mutation rates (Travis and Travis 2002). Indeed, co-occurring species can exhibit a range of mutation rates (Baer et al. 2007), 9 which may also play a role in species differences in the degree of local adaptation and subsequent responses to environmental change (Orr and Unckless 2008). Here, I did not allow explicit evolution of dispersal distance (*σ*), though the comparison of population sizes for my species of differing *σ* provide insight into how dispersal would evolve in this system. In a temporally constant environment, dispersal is maladaptive due to the spatial selective gradient (Balkau and Feldman 1973). However, once temporal change in environment is introduced, greater dispersal can be favored with the functional form of temporal environmental change determining the optimal *σ* (see Blanquart and Gandon 2011 for more detailed analysis). I did not investigate interspecific variation in phenotypic plasticity, which may supplant local adaptation as a response to environmental gradients. As with migration load, if population size is related to the degree of local adaptation versus plasticity (i.e. habitat specialization versus generalization) then changing environments may cause complex community change. Under some models, greater dispersal across environmental gradients can favor plastic responses to environment (Sultan and Spencer 2002, reviewed by Hendry 2016).

The form of environmental change may have dramatic effects on how eco-evolutionary responses influence communities. Previous theory has shown how the benefits of genetic variation (Lande and Shannon 1996) and dispersal (Blanquart and Gandon 2011) can depend on the functional form of environmental change. I found that communities can exhibit distinct dynamics depending on a scenario of abrupt environmental change (Gomulkiewicz and Holt 1995, Orr and Unckless 2008) versus sustained change (Pease et al. 1989, Lynch and Lande 1993, Polechová et al. 2009). Specifically, sustained change favors intermediate gene flow species and results in their stable dominance (highest *N*) in communities, whereas abrupt environmental change results in only transient community change favoring high to intermediate *σ* species. In nature any form is possible and thus my results demonstrate how diverse forms of environmental change may cause complex dynamics in nature.

Though I modeled community turnover in a single local population, all communities in my model are equivalent and the processes I described would occur across species ranges. This suggests that there is a large potential spatial extent of eco-evolutionary responses to rapid environmental change, resulting in community change across large regions. In nature *b* is non-linear and rugged, a feature worthy of study in future simulation of response to temporal environmental change. Furthermore, multiple traits may be under simultaneous spatially-varying selection (Guillaume 2011, Duputié et al. 2012, MacPherson et al. 2015) and selective regimes on these traits may change simultaneously. Given that environmental change can be complex, with different forms of change in different environmental dimensions, it is possible that in nature changes in selective gradients may take multiple functional forms simultaneously leading to complex changes in relative abundance for species differing in *σ*.

The model studied here was simple and thus it is challenging to determine how important my results are in natural systems. However, gene flow across spatial selective gradients is likely a major source of within-population genetic variation in traits under selection (Yeaman and Jarvis 2006, Paul et al. 2011, Farkas et al. 2013). Findings on ponderosa pine suggest that greater *b* can cause greater *V*_*G*_ (Yeaman and Jarvis 2006). Less is known, however, of how adaptability or *V*_*G*_ are related to interspecific variation in population size. The negative relationship between these two quantities is the key to community turnover following environmental change in my results. One problem with empirically studying the processes I described there is often a substantial lag before better dispersing species dominate communities (Figures 2 & 5). Thus researchers may overlook empirical population changes caused by environmental change.

Strongly interacting species may often experience selective gradients driven by the same environmental variable (e.g. temperature, Aitken and Bemmels 2016) while their diversity is also shaped by interspecific variation in dispersal ability (Lasky et al. 2017). Differences among these species in local adaptation to the same environmental variable might lead to different eco-evolutionary responses to environmental change, causing indirect effects on interacting species (Fussmann and Gonzalez 2013). For example, multiple competing tree species may simultaneously be locally-adapted along environmental gradients (Ikeda et al. 2014). Recent work by Brans et al. (2017) has shown similar intraspecific trait clines in multiple co-occurring cladocerans along urbanization gradients drives community patterns. Here I simulated competing species, but interactions of different types (e.g. trophic) may yield distinct eco-evolutionary community responses to changing environments. The competition simulated here was independent of the trait under changing selection, i.e. there was no density or frequency-dependent selection. This scenario may be appropriate for abiotic stressors such as cold or heat. However, other environmental conditions such as resource supply rates might change optimal trait values, but also be subject to density or frequency-dependent selection. These forms of selection can result in more complex patterns of population and trait variation among species (Roughgarden 1976).

My work demonstrates how interspecific variation in gene flow alters communities experiencing environmental change. Some have suggested assisted gene flow as a technique to mitigate climate change impacts on wild populations, with gene flow facilitating local adaptation of populations suddenly experiencing novel climates (Aitken and Whitlock 2013). My results highlight how such approaches could have important effects on community structure. Aitken and Whitlock (2013) suggested that assisted gene flow efforts should be focused on ecologically dominant species (due to importance for ecosystem functioning) and rare species (to prevent extinction). My results show how such a strategy would likely change community structure, as species not included (historically intermediate abundance species) in assisted gene flow would be expected to decline in abundance due to slower adaptation to climate change. Others have suggested breeding of wild species to promote adaptation to future environments (Oppen et al. 2015). These management efforts would have to be balanced across species of different abundances if they are to limit impacts on community composition and ecosystem function.

## Conclusion

Community composition is defined by the population sizes of component species, but greater population size might not correspond to greater adaptability to environmental change. This discrepancy can result in complex community turnover as selection regimes shift. The simple models studied here demonstrate some of the complexity in eco-evolutionary community change. Future research could improve our ability to predict responses to environmental change in nature by learning more about the genetics and ecology of adaptation in addition to theoretical investigation of more complicated scenarios.

## Acknowledgments

This manuscript benefited from comments by Jitka Polechová and two anonymous reviewers, as well as Hidetoshi Inamine, Martin Turcotte, Andrew Gonzalez, and Andrew Hendry, in addition to conversations with Andrew Read, Katriona Shea, and Timothy Reluga.

## Appendix Impacts on community-mean traits

Interspecific variation in adaptation following environmental change will likely have impacts on the distribution of traits in a community, which is often of interest to community and ecosystem ecologists (Muscarella and Uriarte 2016, Šímová et al. 2018). For example, ecosystem function may be influenced by the mass-averaged functional traits in a community (Grime 1998). Under the scenario of abrupt environmental change, the slow adaptation and return to equillibrium abundance of species that 6 dominate communities may have interesting effects on changes in community-weighted mean (CWM) traits. Indeed, following abrupt environmental change, initially there is a very rapid phase of change in CWM driven by fast-adapting high *σ* species (Figure S1). However, there is an abrupt slow-down in change in CWM as most 20 high *σ* species have adapted but low *σ* species remain maladapted. Nevertheless, the 21 low migration load of these low *σ* species contributes to their fitness and abundance and hence influence over CWM traits. When interspecific competition occurs, the change in CWM is even faster because slow-adapting species are supressed (Figure S1). By contrast, when there is sustained change in *θ* over time, species exhibit marked variation in their ability to adapt to the moving optimum. Although the highest *σ* species are able to maintain *z̄* close to the optimum, they are less abundant than intermediate *σ* species due to migration load (equation 10, Figure 6). Thus the CWM exhibits a substantial and stable lag behind the optimum.

## Relationship of eco-evolutionary community turnover to ecological succession

The transient dominance of species with higher gene flow following an abrupt environmental change is qualitatively similar to classic hypotheses explaining successional turnover in communities. Specifically, early successional species may have better dispersed propagules but lower fitness compared to later successional species. In the present model, gene flow and propagule dispersal are one in the same (*σ*), i.e. there is no mechanism of gene flow apart from propagule movement (no gamete movement). To more formally investigate the similarity with succession, I studied how species differing in *σ* in the present model respond to ecological disturbance, with no change in *θ*. In the absence of any environmental change, consider an ecological disturbance that reduces locally-adapted populations of different species by the same large proportion. For simplicity, I assumed a localized disturbance that introduced non-zero 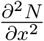(equation 3) but did so orthogonally to *b* such that asymmetric migration had no effect on trait evolution (i.e. 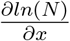) set equal to zero in equation 2).

**Figure 1:**
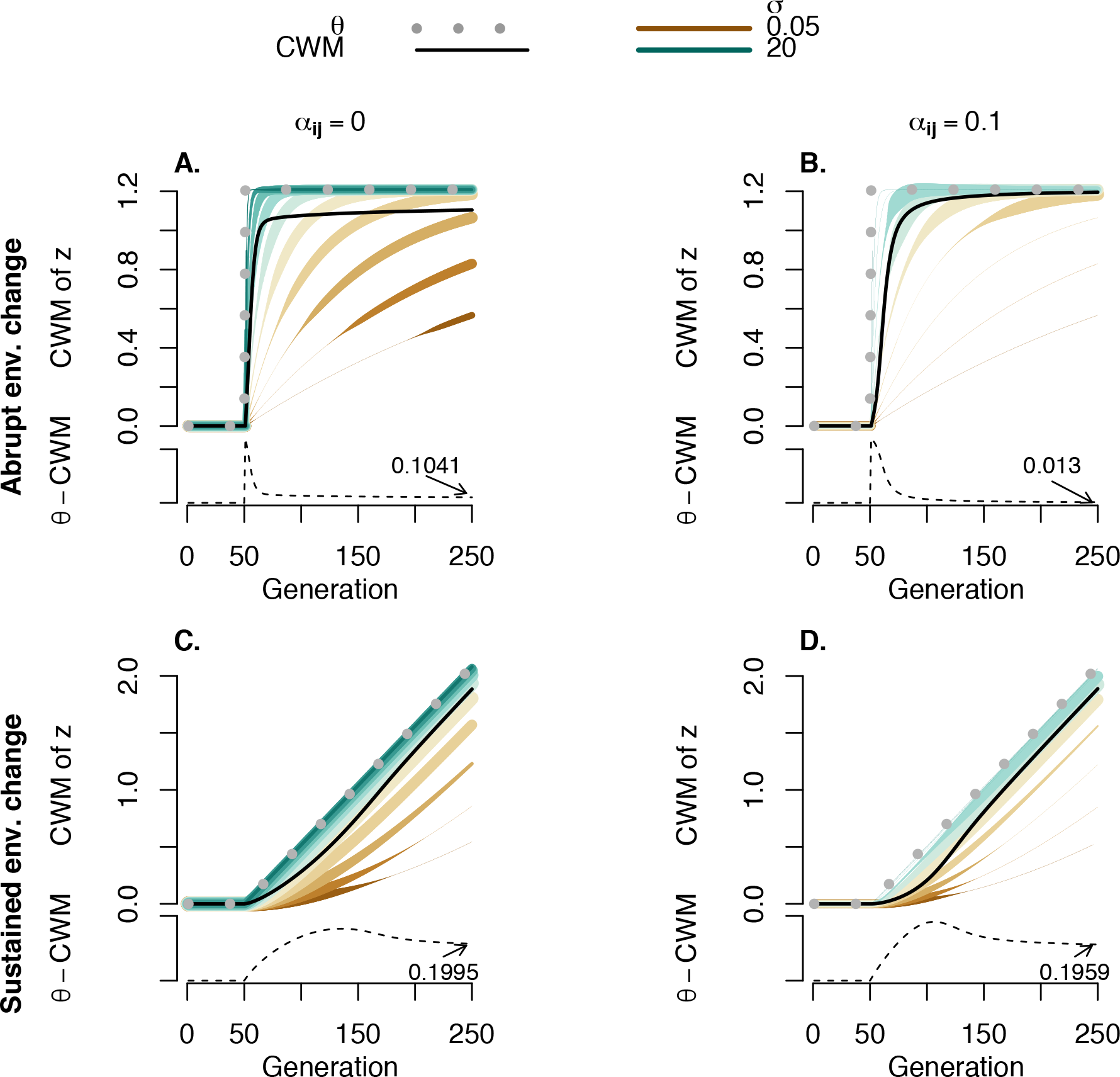
Effects of environmental change on community-weighted mean (CWM) traits under selection due to eco-evolutionary responses. Example species with a range of *σ* values are shown (colors), with line thickness indicating relative abundance (thickness scales not consistent across panels). As in earlier presented simulations, communities were composed of species with a log uniform distribution of *σ* values. The CWM (black line) at each timepoint is an abundance-weighted average of *z*. Small subpanels show the value of the lag in CWM behind *θ* with the final value givien in text. Parameter values (unless otherwise noted) were *b* = 0.05, *V*_*S*_ = 1, *V*_*E*_ = 0.05, and *r*_*m*_ = 0.5. For (A-B), *θ*\* − *θ* = 1.21. For (C-D), *k* = 0.01.

**Figure 2:**
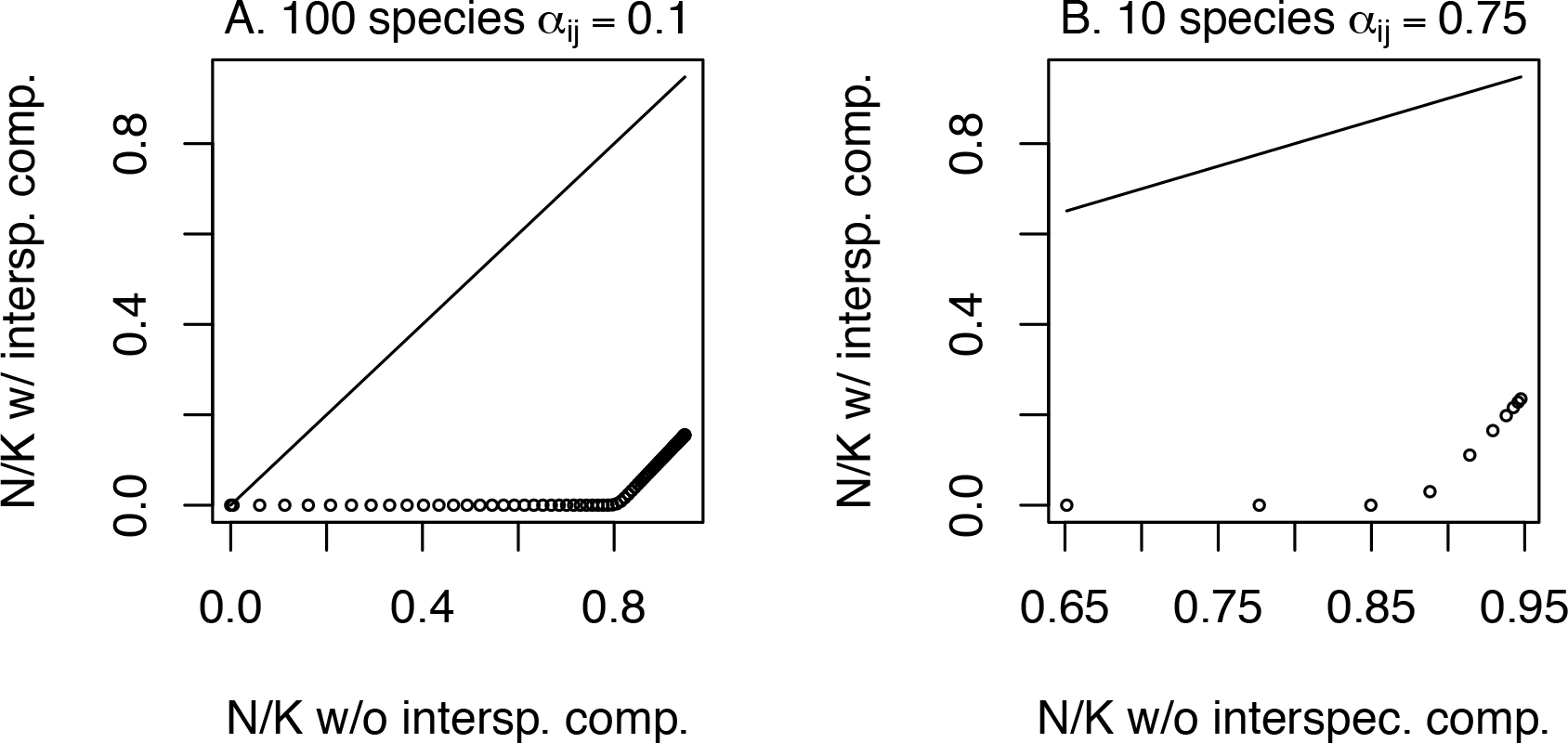
Comparing approximate equilibrium species’ *N* for scenarios differing in presence of interspecific competition. (A) Black circles show species in a diverse (*J* = 100 species) community with weak interactions (*α*_*ij*_ = 0.1). Straight line shows one-to-one relationship. (B) Here the trend in abundance for another (*J* = 10 species) community is shown, where species strongly compete (*α*_*ij*_ = 0.75) according to Lotka-Volterra models. Parameter values (unless otherwise noted) were *b* = 0.05, *V*_*S*_ = 1, *V*_*E*_ = 0.05, and *r*_*m*_ = 0.5.

**Figure 3:**
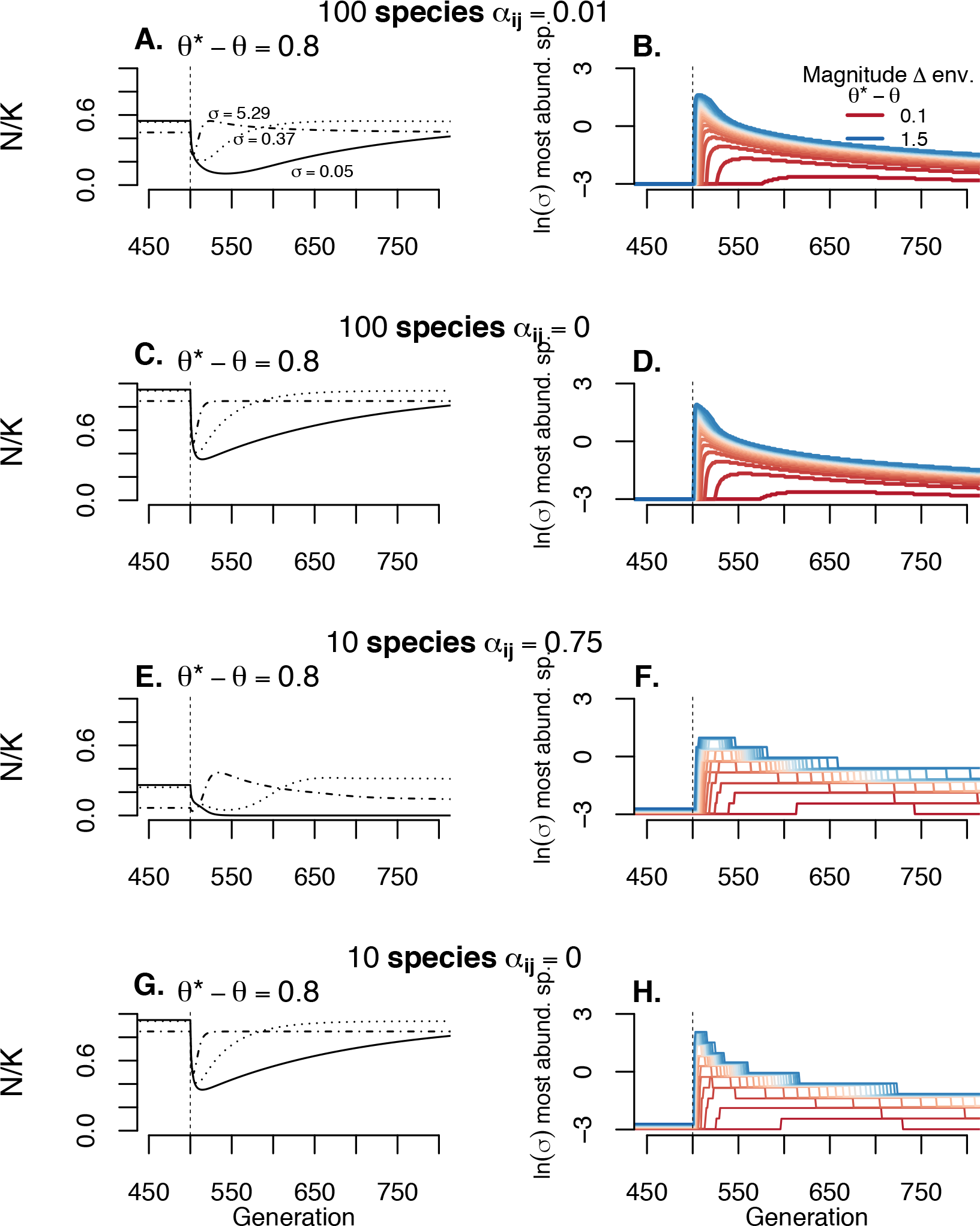
How the magnitude of environmental shift and interspecific competition affects community turnover. Left panels show the same three representative species of varying *σ* under different scenarios of interspecific competition. Right panels show which species are most abundant at any point in time, under different levels of abrupt environmental change. Populations are at approximate equilibrium and adapted to *θ* for the first 500 generations, when an instantaneous environmental change to *θ*\* occurs. Parameter values (unless otherwise noted) were *b* = 0.05, *V*_*S*_ = 1, *V*_*E*_ = 0.05, and *r*_*m*_ = 0.5.

Successional community turnover arises as species differ in the rate of population growth (eqn 3) due to interspecific variation in immigration (favoring high *σ* species) and fitness (favoring low *σ* species). However, note that the fitness advantage of low *σ* species is dependent on reproduction by individuals already present, which are few after disturbance. My simulations showed that the more intense the disturbance, the slower the return to community equillibrium (Figure S4), analogous to the slower return following greater abrupt changes in *θ* (Figure 2). Under a sustained ecological disturbance (constant proportion of individuals lost each generation) ecological community turnover exhibits qualitatively similar patterns to the eco-evolutionary response to sustained change in *θ* (Figure S4). Specifically, sustained disturbance resulted in consistent dominance by species with intermediate *σ*, similar to these species being most abundant under sustained change in *θ* (Figure 4).

**Figure 4:**
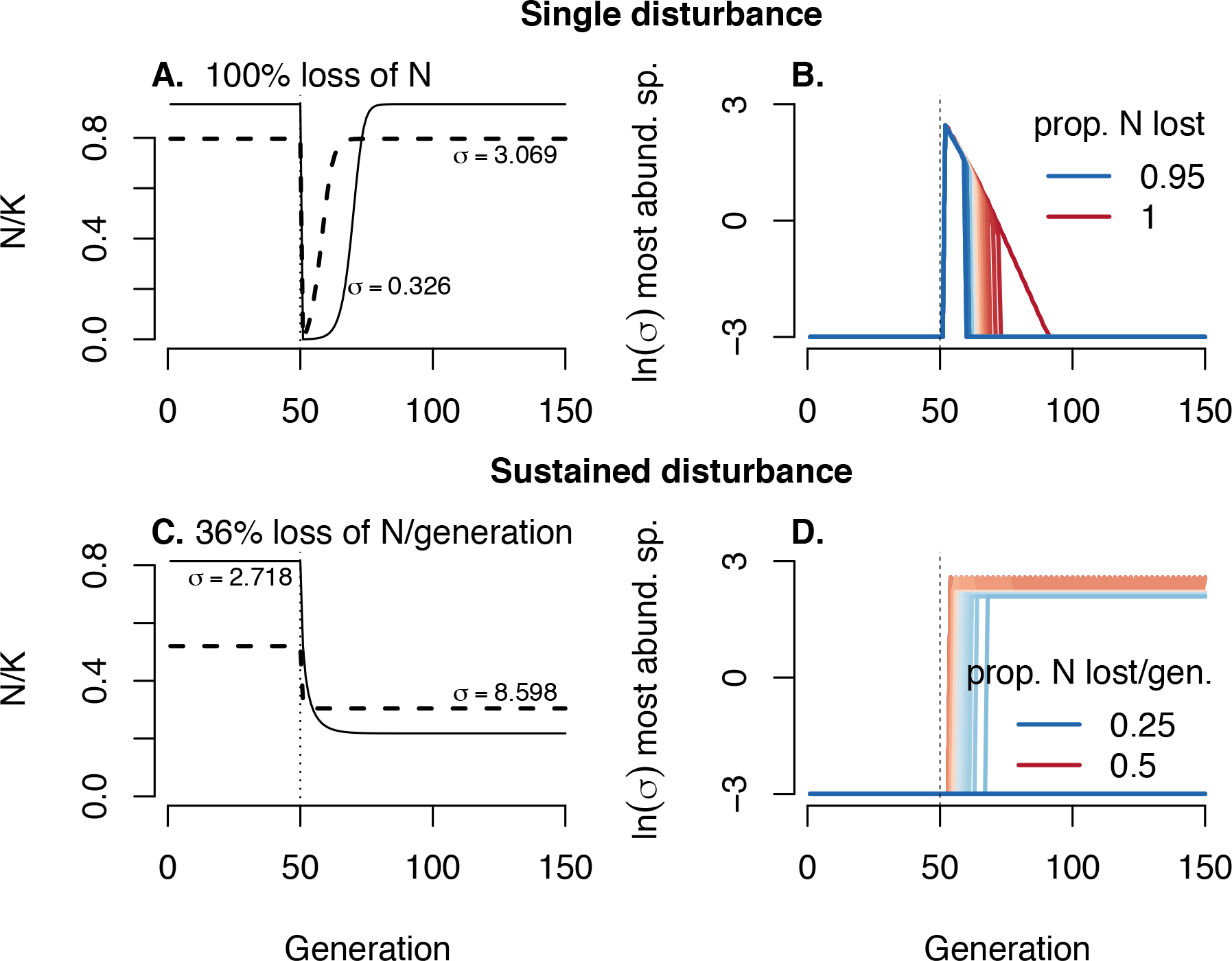
Variation in dispersal (*σ*) among non-interacting species (*alpha*[*ij*] == 0) determines how communities of locally-adapted populations respond to ecological disturbance. (A-B) A single disturbance removes a large portion of each species’ *N* after generation 50. (C-D) recurring disturbances are imposed in each generation, starting after generation 50. Parameter values (unless otherwise noted) were *b* = 0.05, *V*_*S*_ = 1, *V*_*E*_ = 0.05, *r*_*m*_ = 0.5, and 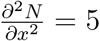.

**Figure 5:**
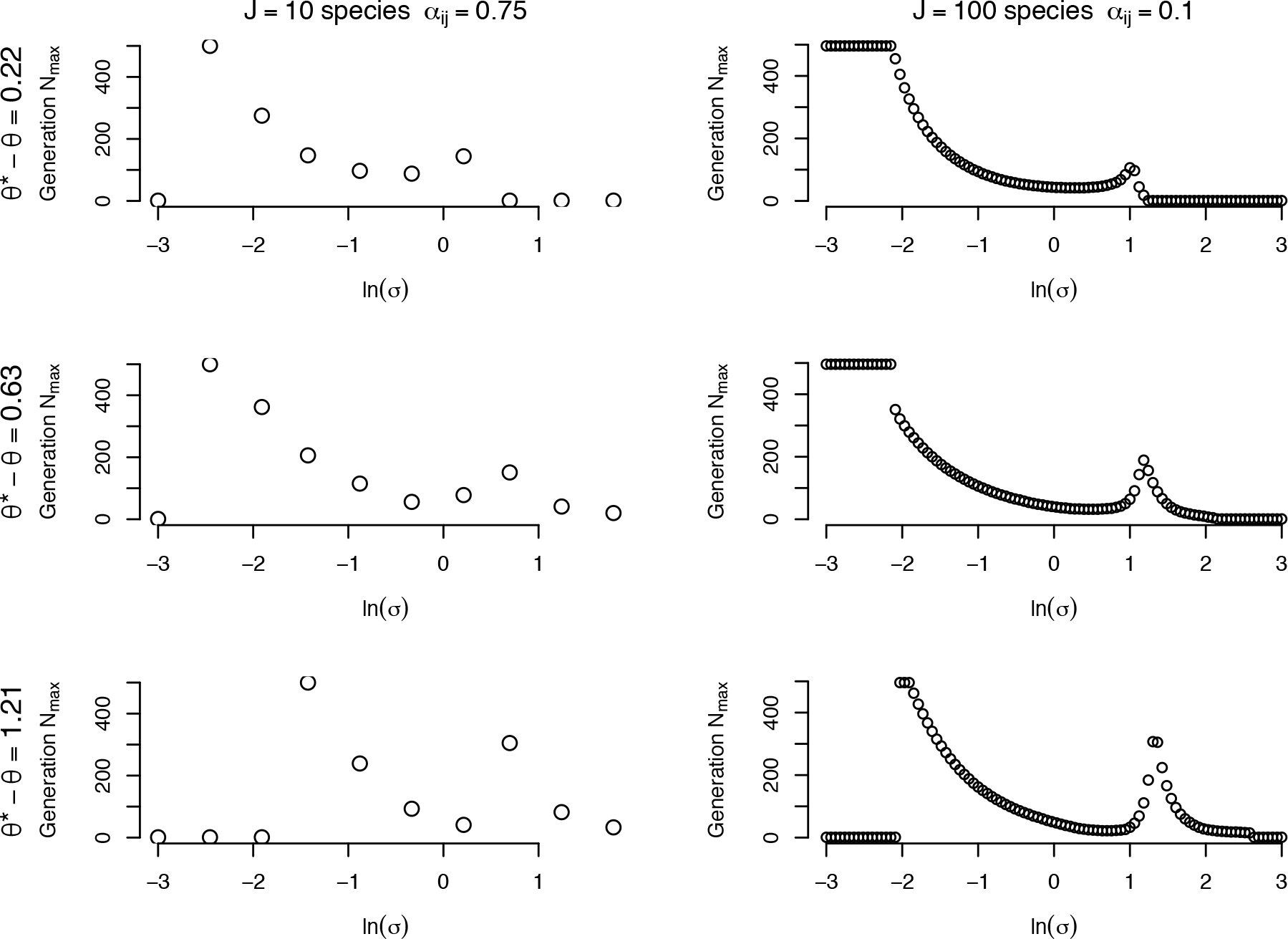
Species that differ in *σ* show different timing of their peak population size (*N*_*max*_) after environmental change. Each panel shows a different scenario of abrupt environmental change. Generation 0 is the generation when environmental change occurs. When there is no interspecific competition (*α*_*ij*_ = 0) species are always most abundant before environmental change (generation 0 on y-axes here). Parameter values (unless otherwise noted) were *b* = 0.05, *V*_*S*_ = 1, *V*_*E*_ = 0.05, and *r*_*m*_ = 0.5.

**Figure 6:**
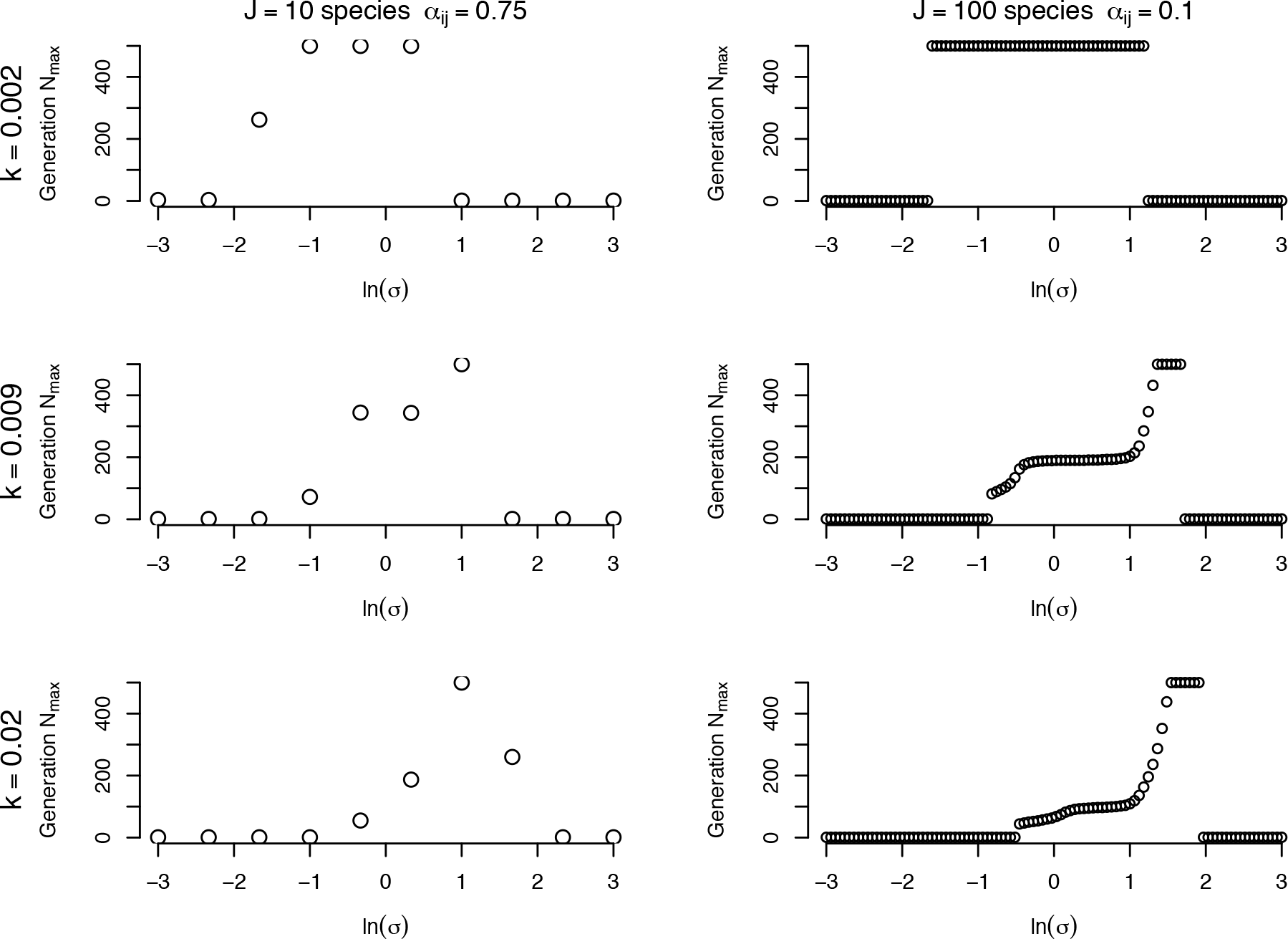
Species that differ in *σ* show different timing of their peak population size (*N_max_*) after environmental change. Each panel shows a different scenario of sustained environmental change. Generation 0 is the generation when environmental change begins. When there is no interspecific competition (*α*_*ij*_ = 0) species are always most abundant before environmental change (generation 0 on y-axes here). Parameter values (unless otherwise noted) were *b* = 0.05, *V*_*S*_ = 1, *V*_*E*_ = 0.05, and *r*_*m*_ = 0.5.

